# Social status and previous experience in the group as predictors of long-term welfare of sows housed in large semi-static groups

**DOI:** 10.1101/2020.12.16.423029

**Authors:** Sophie Brajon, Jamie Ahloy-Dallaire, Nicolas Devillers, Frédéric Guay

## Abstract

Mixing gestating sows implies hierarchy formation and has detrimental consequences on welfare. The effects of social stress on the most vulnerable individuals may be underestimated and it is therefore important to evaluate welfare between individuals within groups. This study aimed at investigating the impact of social status and previous experience in the group on well-being of sows housed in large semi-static groups (20 groups of 46-91 animals). We assessed aggression (d0 (mixing), d2, d27, d29), body lesions (d1, d26, d84) and feeding order. Social status was based on the proportion of fights won during a 6-hr observation period between d0 and d2. Dominants (29%) were those who won more fights than they lost, Subdominants (25%) won fewer fights than they lost, Losers (23%) never won any fight in which they were involved while Avoiders (23%) were never involved in fights. Resident sows (70%) were already present in the group in the previous gestation while New sows (30%) were newly introduced at mixing. Subdominants and Dominants were highly involved in fights around mixing but this was more detrimental for Subdominants than Dominants, Losers and Avoiders since they had the highest body lesion scores at mixing. Avoiders received less non-reciprocal agonistic acts than Losers on d2 (*P*=0.0001) and had the lowest body lesion scores after mixing. However, Avoiders and Losers were more at risk in the long-term since they had the highest body lesions scores at d26 and d84. They were followed by Subdominants and then Dominants. New sows fought more (*P*<0.0001), tended to be involved in longer fights (*P*=0.075) around mixing and had more body lesions throughout gestation than Resident sows. Feeding order from one-month post-mixing was influenced both by the previous experience in the group and social status (*P*<0.0001). New sows, especially with a low social status, are more vulnerable throughout gestation and could serve as indicators of non-optimal conditions.

## Introduction

Due to considerable changes in the pig industry worldwide in favour to groups-housing systems, understanding social behaviour of pigs and its impact on welfare has become increasingly important. The wild counterparts of pigs are highly gregarious and form complex hierarchical structures of multigenerational and matrilineal social units centered around several philopatric females associated with their cohorts of offspring (1,2). Despite occasional breakdown in social cohesion, feral pigs form strong social bonds with related individuals and aggression is very limited (3). Although the domestication process resulted in profound morphological and reproductive changes in pigs, social behaviours were highly preserved and pigs still need to establish a social hierarchy to avoid future conflicts and have good social cohesion (4,5). In the line with this, recent studies provided evidence in growing pigs that high levels of aggression around mixing predicted better future social cohesion and prevented chronic aggression (6). However, under commercial conditions, repeated mixing of unacquainted pigs of similar age and body size causes repeated hierarchy breakdown, disrupts social relationships and results in aggressive contests between group-members within 48 h post-mixing (7), but also in non-reciprocal agonistic interactions (mostly limited to single bites, knocks and threats) on the long-term (8). This unstable social environment can cause stress and injuries (9), impair immune function (10), decrease sows’ productivity (11), negatively modulate maternal behaviour (12) and have negative effects on offspring welfare (13) and daughters’ future maternal behaviour (14) through prenatal stress.

Although welfare state is defined at the individual level and depends on how the animal perceives its environment (15,16), sow aggression and welfare is rarely assessed at the individual level, meaning that the stress and injuries of the most vulnerable individuals may be underestimated (17). Yet, the welfare level of individual sows within a social group may differ according to the social status of the animal (18,19). Findings around mixing are contradictory in gestating sows and, for example, some authors reported no relationship between social status and body lesions (20), more body lesions in low-ranking sows (21) or more body lesions in aggressive sows which were considered as high-ranking sows (22). Conflicting results between studies might be partly explained by housing or social condition differences (e.g., group size). However, low-ranking sows appear to be more vulnerable to social stress and have a greater risk of poor welfare on the long-term. Indeed, previous studies indicated that low-ranking sows had lower risk to develop stereotypies but they spent less time in resting areas (i.e., suggesting that their access to preferred resting areas was denied), they fed later than higher-ranking sows and they showed more body lesions throughout gestation than high-ranking sows (20,23–25). Investigating sow welfare within groups is also important because low-ranking sows were reported to have a significantly lower body weight, farrowing rate, and total litter size, and their offspring had lower weight gain and lean tissue (21,26–28).

In a study with gestating sows mixed into groups of 10 individuals, Verdon et al. (17) found that subdominants (i.e., that received more aggression than they delivered) represented the majority of the groups (44.2-45.5%) while subordinates (i.e., delivered very little or no aggression) were the least common (19.0-22.6%). However, Samarakone and Gonyou (29) suggested that domestic growing pigs may adopt less aggressive social strategies in large groups, with several individuals able to avoid fighting, probably because the risk to lose a fight and the cost of engaging in fights and being severely injured may be higher in large groups where opportunities to fights are more frequent than in small groups. Therefore, the proportion of dominant, subdominant and subordinate sows within a group may differ with group size, along with the impact of social status on welfare.

Most of the previous studies were performed with small group sizes (e.g., 15 individuals or less) while social strategies differ according to the group size (30,31), and groups can count many more individuals (e.g., 50-100 individuals) in commercial farms. Hence, this study aimed at (1) determining the social status of gestating sows housed in a semi-static large group in a commercial setting (up to 91 animals/group) and (2) investigating the impact of social status on welfare indicators related to social behaviour (i.e., agonistic behaviour, body lesions and feeding order) and physiological functioning (i.e., individual and reproductive performance). The influence of a previous experience in the group (i.e., Resident or New) on welfare was also considered in the analyses. While agonistic behaviour was recorded at mixing and one month later, body lesions were measured the day after mixing, as well as one- and three-months post-mixing. It was hypothesised that low-ranking sows (i.e., Losers and Avoiders), which win no fights on the day of mixing, would represent a higher proportion of the group than in previous studies with small group sizes (e.g., Verdon et al., 2016) but would still be the principal recipients of aggression and suffer from more injuries later during gestation, and feed later than Dominant and Subdominant sows. In addition, New sows may suffer from more aggression and body lesions around mixing, compared to Resident sows who were already part of the groups during the previous gestation, but may not differ later during gestation once the hierarchy is formed.

## Material and Method

The trial was performed in a gestation unit of a commercial 900 sows breed-to-wean farm located in Chaudière-Appalaches (Quebec, Canada) between February 2019 and February 2020. An initial paper recently published (32) investigated sows’ welfare at the group level and showed that sows’ behaviour and welfare differed according to the genetic line. In contrast, the present study aimed at exploring the intra-group variability in welfare at the individual level. The 900 sows were distributed between five cohorts of two groups of up to 91 animals. Sows within groups were synchronized so that two groups were inseminated every 28 days and so on. Lighting was provided from 0700 to 1900. Ventilation fans and heaters switched on automatically according to the temperature so that the mean daily ambient temperature was maintained between 19.2-22.2°C throughout the year.

This study was carried out in accordance with the recommendations in the “Canadian Code of practice for the care and handling of pigs” (33). All procedures on animals were conducted after prior approval of the Laval University Institutional animal care committee (Quebec city, Quebec, Canada, approval number: 2018102-1). Animals’ health was monitored daily throughout the experiment and sows were removed from the group if showing moderate to severe lameness or suffering from significant injury or illness.

### Animals and housing

This study comprised 646 multiparous gestating sows of mixed parities (range 1-5). Single high performance genetic lines (HP1 or HP2 genetic line, both F1 derived from Landrace x Yorkshire sows and developed by two different breeding companies) groups of 44 to 91 animals were studied across one to three gestations (between the 3^rd^ and 5^th^ rotation) for a total of 10 replicates including one group of each genetic line (i.e., total of 20 groups). The rotation number corresponded to the insemination number (e.g., rotation 3 = third insemination of the cohort) but sows within the cohort could have a lower parity (up to two gestations late) if they had previous reproductive failures and were moved to the next cohort. At 5-7 days post-insemination, sows were moved from their insemination stalls and mixed into one of the six large gestation pens (day of mixing: d0). Each pen was 9.45 m long and 21.34 m wide (total of 201.66 m^2^), which provided at least 2.21 m^2^ floor space allowance per sow. Pens had part slatted concrete floor with four plastic walls in a central area and five solid concrete resting areas (6.89 m^2^ each) separated by concrete walls in the back area. Each pen was equipped with three water bowls, four nipple drinkers and one single-entry/exit Electronic Sow Feeder (ESF, Compident ESF, Schauer Agrotronic GmbH, Prambachkirchen, Austria). The ESF opened at 2200 h and closed once all the sows were fed or one hour after the last visit if one or two sows did not feed. When more than two sows did not feed, the ESF closed at 1800 h. Sows were provided wet gestation diet (with 14.2% min. crude protein, and 4.0% min. crude fat, and 7.5% min. crude fiber as dry matter (DM, 88%) basis) formulated to meet the National Research Council nutritional requirements for gestating sows (34).

Pregnancy diagnosis was performed using two methods during the first weeks of gestation. Sows which showed behavioural signs of estrus during the first two weeks after mixing were directly removed from the gestation pen and transferred to the insemination stalls to be re-inseminated and added to the following group of the same genetic line (cohort +1). In addition, pregnancy tests using ultrasound scanning were performed at d23 post-mixing. All sows that tested negative in the pregnancy test were then removed from the group at d28 post-mixing and housed into individual insemination stalls until the following insemination phase one month later (cohort +2). Sows were thus involved in one (N = 200), two (N = 318), three (N = 106), four (N = 20) or five (N = 2) replicates, for a total of 1244 observations. Group composition was always different across gestations since new sows from previous cohorts were always added at mixing while others were removed. Hence, previous experience in the group was considered and, within groups, sows were considered as Resident if they were already present in the group in the previous gestation, or New if they were newly introduced in the group at mixing (average of 30.42 ± 2.90 % of New sows in groups).

The gestating sows remained in the gestation pen for 102 days after mixing. One week prior to farrowing, sows were moved to individual farrowing crates (1.52 m x 2.13 m) where they remained until weaning. At this stage, sows were fed with a lactation diet including 22.0% min. crude protein, 5.3% min. crude fat, and 3.7% min. crude fiber as DM (88%) basis (34) using an individual automatic feeder (Gestal Quattro, Jyga Technologies Inc.©, St-Lambert-de-Lauzon, Québec, Canada).

### Data collection

#### Agonistic behaviour

Animals were video recorded using two digital HD video camera recorders (Sony CX455 Handycam®, Tokyo, Japan) from 0900 to 1600 h on the day of mixing, and from 0700 to 1130 h the days 2, 27 and 28 post-mixing. Post-mixing behavioural analyses were performed as soon as the light turn on at the morning since preliminary observations indicated that sows were more active and susceptible to attack an opponent at this time. The use of non-competitive ESF systems (i.e., offering full protection to the sow at feeding (35)), prevented from exacerbated aggression at feeding. The agonistic behaviour was coded continuously from the video recordings for 4 h post-mixing, and for 2 h from 0700 h at days 2, 27 and 29 post-mixing using the video analysis software The Observer XT 14.2 (Noldus Information Technology, Wageningen, The Netherlands).

Behaviours recorded as non-reciprocal agonistic acts were threats (the perpetrator suddenly stretches the neck toward a recipient and provokes the recipient avoidance or escape without physical contact), bites and knocks (the perpetrator bites or gives a head knock to the recipient) and bullying (the perpetrator gives a series of three or more threats, bites and/or knocks to the recipient). Fight occurrence and duration were also recorded and were defined as a reciprocal act of at least three bites or knocks where each opponent bites or gives a head knock at least once and for a total duration of at least 3 sec (21,36). The aggressive pattern was investigated by classifying sows in four categories according to their propensity to initiate and receive a fight: Sows not involved in fights: sows which were never involved in fights; Systematically aggressed sows: sows which never initiated a fight but received at least one fight; Intermediate sows: sows which initiated fewer fights than they received; Predominantly aggressor sows: sows which initiated more fights than they received. All the sows were individually identified by colour mark coding on the back on day −1 pre-mixing and day 26 post-mixing during the body lesion scoring (*see below*) in order to be identified. The identity of the initiator (sow which gives the first agonistic act), the receiver (sow which receives the first agonistic act), the winner (sow which gives the last agonistic act) and the loser (sow which avoid or escape after the last agonistic act) were recorded. When there was no clear winner or loser, the fight outcome was considered as undetermined. All video analyses were performed by a single trained experimenter and the percentage of intra-observer agreement was calculated using 20-min video sequences per group for a total of 400 min (intra-observer agreement: sow identity: 91.6 %; behaviour: 89.5 %; fight duration: 94.0 %). Groups were identified by a unique code visible on the cameras allowing the observer to be blind to treatments.

#### Social-rank index calculation

The social-rank index (SRI) was calculated for each individual using the 6-hr fight data at mixing and day 2. The calculation method was adapted from Mendl et al. (1992) but fight outcome (i.e., win or lose), instead of displacement (i.e., displace successfully or not), was used as follow:

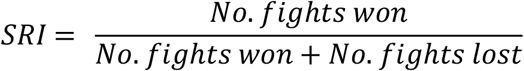

The calculation yielded a SRI score comprised between 0 and 1 which was translated into four social statuses. Sows which won more fights than they lost (SRI > 0.5) were classified as Dominants, sows which won fewer fights or an equal number than they won (0 > SRI ≥ 0.5) were classified as Subdominants, sows which did not win any fight (SRI = 0) were classified as Subordinates. Thereafter, Subordinate sows were classified in two sub-categories: Subordinate which lost all fights were classified as Losers while Subordinate which did not receive any fights were classified as Avoiders.

#### Body lesions

Sow’s body lesions were recorded the day before mixing (day −1 pre-mixing), the day after mixing (day 1 post-mixing), and one (day 26) and three (day 84) months post-mixing. The first body lesion scoring (day −1 pre-mixing) served as the baseline and was subtracted from the body lesion scoring at day 1 post-mixing. The method used to score body lesions was adapted from Turner et al. (37) and Calderón Díaz et al. (38). Three body regions were examined on the left and right sides for a total of six scores per scoring day: front (head, neck, shoulders, and front leg), middle (back and flank), and rear (rump and hind leg). Body lesions for each region and side were scored as follow: 0 = None or 1 superficial lesions; 1 = 2 to 5 superficial lesions; 2 = more than 5 superficial lesions and/or 1 slightly deep red lesions of 2 to 5 cm length; 3 = 2 to 5 slightly deep red lesions of 2 to 5 cm length and/or 1 very deep red lesion of more than 5 cm length; 4 = more than 5 slightly deep red lesions of 2 to 5 cm length and/or 2 to 5 very deep red lesions of more than 5 cm length; 5 = more than 5 very deep red lesions of more than 5 cm length. The body lesion scoring at day 84 post-mixing of the last replicate that had to be recorded in March 2020 was missing due to prohibited access to the farm, as a safety measure, during the worldwide COVID-19 epidemic. The body lesion scores were recorded by a single trained experimenter and the intra-observer agreement was evaluated using body lesion scoring from 304 sows randomly selected at different points of time of the experiment which were scored twice at the same scoring day (intra-observer agreement: 93.2%).

#### Feeding order to the ESF station

The order in which each sow went through the ESF was automatically recorded by the ESF monitoring software (FarmManager software, Schauer®, Austria). Daily data was extracted on average for 13 days (one data/sow/day) spread over the gestation, except the last replicate for which data was extracted for 6 days only due to prohibited access to the farm later during the gestation. Horback and Parsons (22) recorded Feeding Order (FO) from two-weeks post-mixing since they observed that FO was highly unstable during the week of mixing. Preliminary investigation showed comparable inconsistencies between FO in the first days after mixing and FO later during gestation. Thus, the recording days included days one-month (between d24 and d30), two-months (between d52 and 58) and three-months (between d80 and 86) post-mixing. A FO value close to 0 % indicated that the sow went through the ESF early and FO value close to 100 % indicated that the sow was late in the queue and visited the ESF for the first time of the day after the pen mates.

#### Individual and reproductive performance

Measures of backfat thickness before insemination (N_obs_ = 204) and/or before farrowing (N_obs_ =449), as well as measures of body weight before farrowing (N_obs_ = 550) and/or after weaning (N_obs_ = 532) were taken from sows selected randomly and were used as indicators of the individual performance. The success to farrowing (i.e., sows that successfully give birth to at least one alive piglet, N_obs_ = 1242) as well as measures on offspring (N_obs_ = 592) were also recorded and regarded as indicators of reproductive performance. Measures on offspring included the litter size and weight at birth, the number of piglets born alive, stillborn, mummified, total dead and weaned.

### Statistical Analyses

All analyses were conducted using SAS software (version 9.2; SAS Institute Inc., Cary, North Carolina, USA). Preliminary investigation indicated dissociated effects of social status within groups and genetic lines between groups. The within-groups (in this paper) and between-groups (32) effects were thus examined separately. Data were analysed using the sow within the group as the repeated factor, except performance data where the repeated factor was the sow. For continuous data, normality of model residuals was tested using the Shapiro-Wilk test of normality.

First, descriptive analyses were performed in order to illustrate the distribution of sows between social status. The sows’ distribution between social status was expressed as the average percentage (± SE) of sows for each social status across the 20 groups. Chi square tests of independence were performed to examine the relation between social status at the first and the second replicate in which the sows were involved and the relation between social status and aggressive pattern. Pearson (ESF data) and Spearman’s rank (SRI and behaviour data) correlations were performed between the sows’ first two records to estimate the individual sow phenotypic stability over time. Previous analyses indicated that the social status was not significantly associated with the previous experience in the group (χ^2^ (3, N = 1244) = 4.70, *P* = 0.196). Thus, the previous experience in the group variable was considered as a distinct fixed effect in the models.

The effect of social status on agonistic behaviour was analysed using multivariate linear models. Independent variables considered as fixed effects were social status (Dominant, Subdominant, Loser, Avoider), previous experience in the group (Resident, New), rotation (3, 4, 5) and, when applicable, hour (hour 1, 2, 3, 4) or day (day 0, 2, 27, 29) while the group (1 to 20) was considered as random effect. The variables of previous experience in the group, rotation and observation day or hour were removed from the model if non-significant; Otherwise, all interactions were tested and kept when significant. The observation duration differed according to observation day (i.e., four hours at mixing, two hours at d2, d27, d29), so behaviour frequency and total duration were divided by the number of observed hours to have the behaviour frequency and duration per hour. Group size decreased across time, since no further animals were bought from the sows acquisition, and was confounded with rotation and sows’ parity that increased across time. Hence, the rotation variable was taken into account in the analyses and was regarded as an integrative variable taking into account parity and group size. Because behaviour data distributions were skewed, the number of agonistic acts was analysed using the GLIMMIX procedure with a Poisson distribution and log link function while the fight duration was analysed using the GLIMMIX procedure with a lognormal distribution and the identity link function. The denominator degree of freedom of these latter analyses were computed using Satterthwaite approximation. The estimates were then back-transformed to the original scale and model results were expressed as predictive means [CI].

Body lesion scores from the left and right sides of the body were pooled and averaged to keep only one score for the front, middle and rear of the body. In addition, the six body lesion scores were pooled and averaged to create the variable total body lesions. Thereafter, the body lesion scores were re-categorised for each body region in four balanced categories to be properly analysed (very low, low, high, very high). Those ordinal categorical variables were then modelled using multinomial models to fit cumulative logit proportional odds model to the data and odds ratio were calculated. The same independent variables as for the behavioural analyses were considered.

Pearson correlations were performed between the median FO one-, two- and three-months post-mixing to evaluate their consistency across time. Because there was a strong FO consistency between one-, two-, and three-months post-mixing (*see results below*), the median FO between one- and three-months post-mixing was selected for each sow for the subsequent analyses. This variable was then categorised in three balanced categories: FO1 < 34%; 34% ≤ FO2 < 64%; FO3 ≥ 64%. A sow was categorised as FO1 if she was among the first 34% to feed at the ESF. The Cochran-Mantel-Haenszel test (CMH) was used to test the association between FO and social status as well as the association between FO and previous experience in the group.

Individual and reproductive performance were analysed by multivariate linear models with social status, previous experience in the group and rotation considered as fixed effects and replicate considered as random effect. The variables of previous experience in the group and rotation were removed from the model if non-significant; Otherwise, all interactions were tested and kept when significant. Residuals from most of the performance analyses had a normal distribution except the number of piglets stillborn which was modelled with a Poisson distribution and the total number of dead piglets which was modelled with a lognormal distribution. The variables success to farrowing and presence of mummified piglets were binary and were modelled using mixed logistic regression. Model results were expressed as predictive means ± standard errors. All multiple comparisons were performed using Tukey’s adjustment of the Student’s *t* tests.

## Results

### Dominance and consistency of behaviours over replicates

Social groups were characterized by the presence of a high proportion of Subordinate sows (i.e., that did not win any fights) since they represented an average of 45.7 ± 1.5 % of the animals within the group. More particularly, the two sub-categories, Losers (i.e., that lost at least one fight) and Avoiders (i.e., that were not involved in any fight) represented 22.8 ± 1.1 % and 22.8 ± 1.4 % of the animals, respectively. The Dominant sows (i.e., that won more fights than they lost) represented 29.0 ± 1.3 % of the sows while Subdominant sows (i.e., that won as much as or fewer fights than they lost) represented 25.4 ± 1.4 % of the sows within the groups. Spearman’s rank correlations indicated that there was a weak relationship between SRIs over the first two replicates in sows that were involved in fights (N = 308; p = 0.17; *ρ* = 0.003). Avoiders were not considered in those correlations since they were not involved in fights and did not have SRI. Similarly, the sows’ social status in the first replicate in which they were observed was consistent with the social status in the subsequent replicate (χ^2^ (9, N = 446) = 131.3, *P* < 0.0001).

Although the aggressive pattern (i.e., the extent to which a sow initiated a fight) was associated with the social status (χ^2^ (4, N = 947) = 133.4, *P* < 0.0001), the **Table 1** shows that the most aggressive sows were not necessarily the Dominants. For instance, one third of the Losers (7.2 % over the 22.8 %) were predominantly aggressors (i.e., initiated more than half of the fights in which they were involved) but never won a single fight. The aggressive pattern may reflect the temperament of the individuals since there were a consistency of agonistic behaviours at mixing over replicates (Spearman’s rank correlation between the first two replicates where the sow was present: N = 446; 0.42 < ρ < 0.50; *ρ* < 0.0001).

**Table 1.** Proportion of sows exhibiting each aggressive pattern* according to the social status

| Aggressive pattern* | Social Status |  |  |  | Total |
| --- | --- | --- | --- | --- | --- |
|  | Avoider | Loser | Subdominant | Dominant |  |
| Not involved in fights | 22.8 ± 1.4 | - | - | - | **22.8 ± 1.4** |
| Systematically aggressed | - | 11.8 ± 1.2 | 3.4 ± 0.6 | 5.6 ± 0.9 | **20.8 ± 1.8** |
| Intermediate | - | 3.8 ± 0.5 | 11.6 ± 1.1 | 11.5 ± 1.1 | **26.9 ± 1.7** |
| Predominantly aggressors | - | 7.2 ± 0.7 | 10.4 ± 0.6 | 11.9 ± 0.8 | **29.4 ± 0.9** |
| Total | **22.8 ± 1.4** | **22.8 ± 1.1** | **25.4 ± 1.4** | **29.0 ± 1.3** | **100.0** |

\*The aggressive pattern was divided in four categories; Not involved in fights: The sows were not involved in any fight during the observation period; Systematically aggressed : The sows never initiated a fight but received at least one fight; Intermediate : The sows initiated fewer fights than they received; Predominantly aggressors : The sows initiated more fights than they received.

### Agonistic behaviour

#### Agonistic behaviour across hours on mixing day

A detailed analysis was performed on the day of mixing to investigate whether behavioural differences between social statuses were stable across the first four hours post-mixing. **Fig 1** illustrates the influence of social status on the delivery and reception of non-reciprocal agonistic acts (i.e., bites and knocks, threats, bullying) across the first four hours post-mixing. While the number of non-reciprocal agonistic acts delivered by Dominants and Subdominants was high during the first hour post-mixing and decreased gradually in the subsequent hours (*F*_3,3710_ = 17.3 and 15.5, *P* < 0.0001 and *P* < 0.0001, respectively), it was low and stable for Losers (*F*_3,3710_ = 2.1, *P* > 0.10). It was also low for Avoiders but slightly increased across time (*F*_3,3710_ =3.5, *P* = 0.010). The proportion of non-reciprocal agonistic acts received by Dominants and Subdominants decreased across time (*F*_3,3710_ = 7.8 and 5.2, *P* < 0.0001 and *P* = 0.001, respectively) while it was stable both for Losers and Avoiders during the first hours after mixing (*F*_3,3710_ = 0.3 and 0.9, *P* > 0.10 and *P* > 0.10, respectively). The previous experience in the group also had an impact on the frequency of non-reciprocal agonistic acts delivered and received in certain observation hours. Indeed, Resident sows delivered more non-reciprocal agonistic acts during the fourth hour post-mixing than New sows (1.07 [0.97;1.17] vs. 0.86 [0.74;0.98], *F*_1,3707_ = 7.4, *P* = 0.006), but not during the three first hours post-mixing (*P* > 0.10). In addition, Resident sows received more non-reciprocal agonistic acts than New sows during the first two hours post-mixing (h1: 2.21 [2.09;2.34] vs. 1.81 [1.66;1.98], *F*_1,3707_ = 13.9, *P* = 0.0002; h1: 2.00 [1.89;2.12] vs. 1.70 [1.55;1.86], *F*_1,3707_ = 9.0, *P* = 0.003).

**Fig 1.**
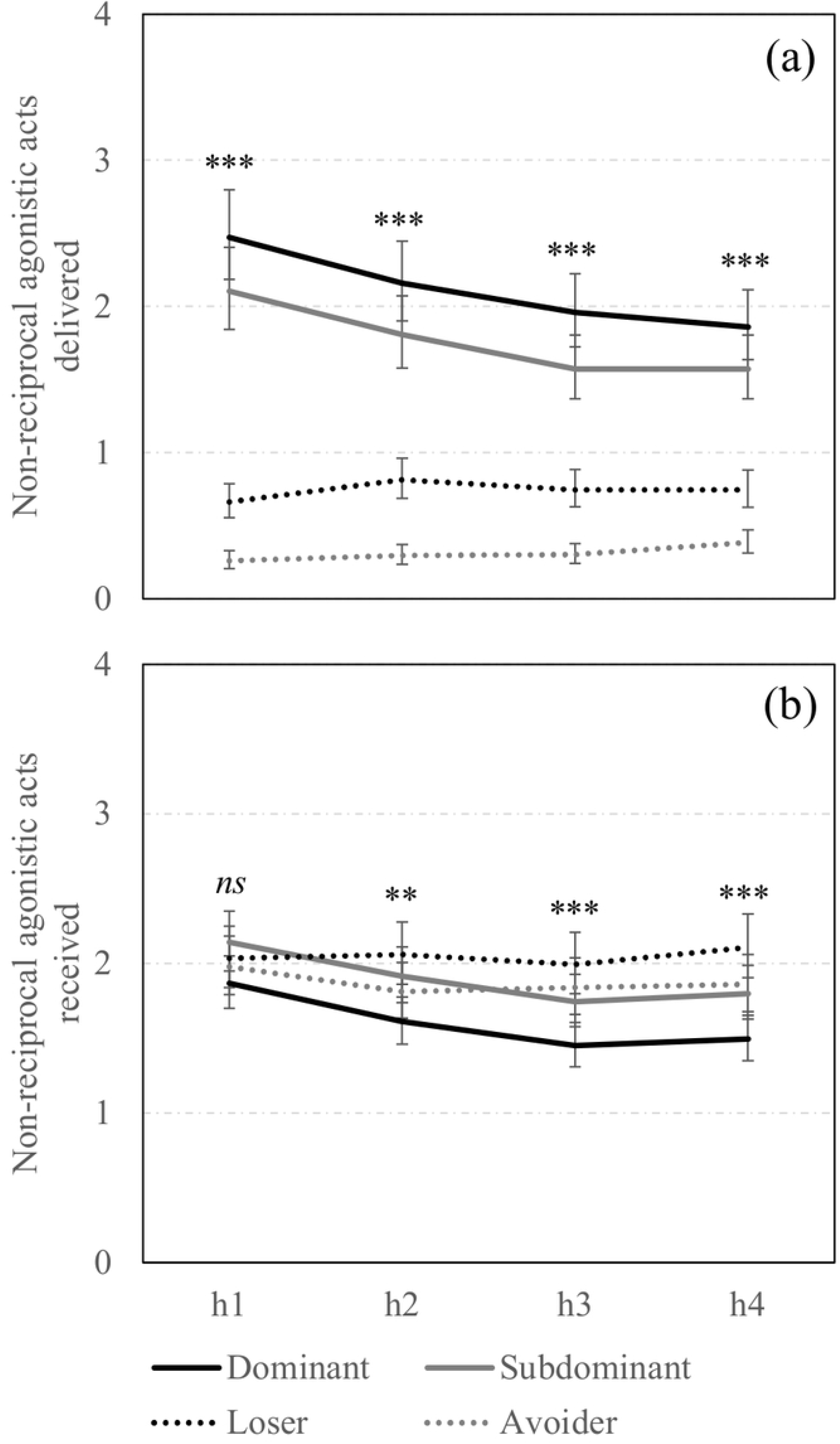
Non-reciprocal agonistic acts across hours on mixing day according to social status. Back-transformed predicted mean [CI] of total non-reciprocal agonistic acts (i.e., bites and knocks, threats, bullying) (a) delivered and (b) received on mixing day across observation hours (h1, h2, h3, h4) and according to the social status (Dominant: solid black line; Subdominant: solid grey line; Loser: dotted black line; Avoider: dotted grey line).**,***; Means differ significantly (** *P*<0.010, *** *P*<0.001); † Means tend to differ (*P*<0.10) between social statuses for a same observation day.

The mean number of fights in which sows were involved was particularly high during the first hour post-mixing, and then decreased by two thirds only two hours later (1.66 [1.57;1.75], 0.72 [0.67;0.78], 0.55 [0.50;0.60], 0.53 [0.48;0.57] fights during the first, second, third and fourth hour post-mixing, respectively, *F*_3,3753_ = 313.9, *P* < 0.0001). Social status also had an impact on fight frequency at mixing since Dominants and Subdominants were involved in an average of 1.10 [1.04;1.15], and 1.07 [1.01;1.13] fights, respectively, while Losers were involved in 0.39 [0.35;0.42] fights (*F*_2,3753_ = 214.5, *P* < 0.0001), independently of the observation hour. New sows were involved in 0.88 [0.83;0.94] and Resident sows in 0.67 [0.63;0.70] fights per hour on mixing day (*F*_1,3753_ = 64.9, *P* < 0.0001), independently of the observation hours.

Taken together, high-ranking sows were engaged in more agonistic acts (i.e., total number of reciprocal and non-reciprocal agonistic acts delivered and received) than low-ranking sows on the day of mixing, however, there was a significant social status x observation hour interaction effect (*F*_9,3710_ = 12.5, *P* < 0.0001). Indeed, Dominant and Subdominant sows were particularly involved in aggression at mixing and their involvement decreased across time (Dominants: 6.91 [6.45;7.40], 5.14 [4.79;5.53], 4.47 [4.16;4.82], 4.44 [4.12;4.78] agonistic acts during the first, second, third and fourth hour post-mixing, respectively, *F*_3,3710_ = 103.3, *P* < 0.0001; Subdominants: 6.71 [6.24;7.22], 5.00 [4.63;5.39], 4.35 [4.02;4.70], 4.42 [4.09;4.78] agonistic acts during the first, second, third and fourth hour post-mixing, respectively, *F*_3,3710_ = 82.6, *P* < 0.0001). In contrast, Loser and Avoider sows maintained relatively low levels of aggression and were less involved in aggression across the first four hours after mixing (Losers: 3.55 [3.25;3.87], 3.39 [3.10;3.71], 3.13 [2.86;3.43], 3.30 [3.02;3.61] agonistic acts during the first, second, third and fourth hour, respectively, *F*_3,3710_ = 2.8, *P* = 0.038; Avoiders: 2.28 [2.07;2.50], 2.13 [1.94;2.35], 2.10 [1.91;2.32], 2.25 [2.04;2.47] agonistic acts during the first, second, third and fourth hour, respectively, *F*_3,3710_ = 1.1, *P >* 0.10).

#### Agonistic behaviour across observation days

Agonistic behaviour was then analysed across observation days. Because the observation duration varied according to observation days, the mean number of agonistic acts per hour on each observation day was calculated and used for the analyses. The **Fig 2** and **3** summarize the effect of social status on non-reciprocal agonistic behaviours delivered and received per hour by the sows on each observation day and according to their previous experience in the group.

**Fig 2.**
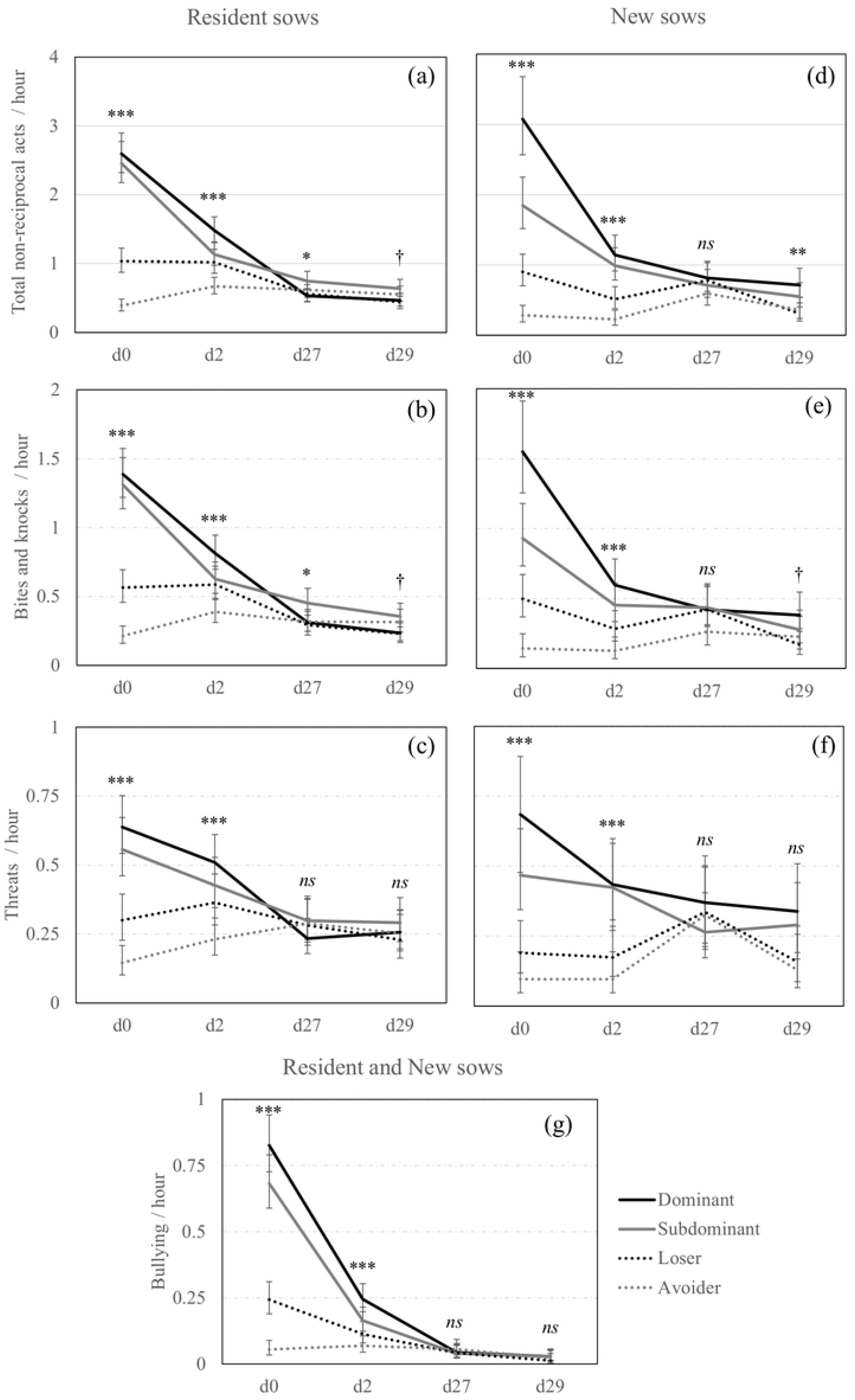
Non-reciprocal agonistic acts delivered across days according to previous experience in the group and social status. Back-transformed predicted mean [CI] number of (a) total non-reciprocal agonistic acts, (b) bites and knocks, and (c) threats delivered per hour by Resident sows, (d) total non-reciprocal agonistic acts, (e) bites and knock, and (f) threats delivered per hour by New sows and (g) bullying delivered per hour by all sows independently of their previous experience in the group, across observation days (d0, d2, d27, d29) and according to the social status (Dominant: solid black line; Subdominant: solid grey line; Loser: dotted black line; Avoider: dotted grey line). *,**,***; Means differ significantly (* *P*<0.050, ** *P*<0.010, *** *P*<0.001); † Means tend to differ (*P*<0.10) between social statuses for a same observation day.

**Fig 3.**
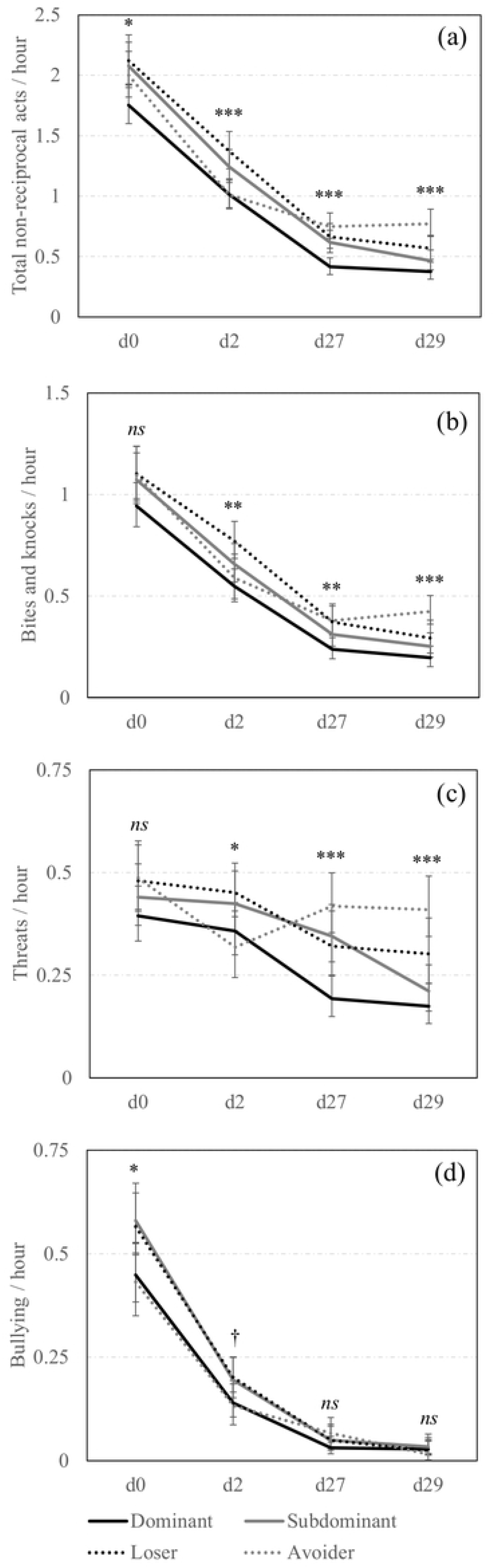
Non-reciprocal agonistic acts received across days according to social status. Back-transformed predicted mean [CI] of number (a) total non-reciprocal agonistic acts, (d) bites and knocks, (c) threats and (d) bullying received per hour by all sows across observation days (d0, d2, d27, d29) and according to the social status (Dominant: solid black line; Subdominant: solid grey line; Loser: dotted black line; Avoider: dotted grey line). *,**,***; Means differ significantly (* *P*<0.050, ** *P*<0.010, *** *P*<0.001); † Means tend to differ (*P*<0.10) between social statuses for a same observation day.

There was a significant triple interaction effect between social status x previous experience in the group x observation day for all the non-reciprocal agonistic acts delivered except bullying (**Fig 2**). At mixing and day 2, among the Resident sows, Dominant and Subdominant sows delivered the numerically highest total amount of non-reciprocal acts (i.e., bites, knocks, threats and bullying pooled together), Losers were the next, and Avoiders were the least aggressive. The same effect was observed between the New sows. Behavioural differences between social statuses were weaker one-month post-mixing, at days 27 and 29 and pairwise comparisons using Tukey’s adjustment were non-significant.

Regarding the recipients of non-reciprocal agonistic acts (**Fig 3**), the social status x observation day interaction was significant for all the non-reciprocal agonistic acts received by the sows except bullying. There was an impact of social status on the reception of non-reciprocal agonistic acts around mixing. Indeed, when considering all the non-reciprocal agonistic behaviours, Loser and Subdominant sows were the most frequently aggressed on day 2 but the pairwise comparisons using Tukey’s adjustment were non-significant on day 0. One month later, on days 27 and 29, sows from low social status still received significantly more non-reciprocal agonistic acts than Dominant sows. Also, the frequency of non-reciprocal agonistic acts received by New sows did not significantly differ from that received by Resident sows.

Reciprocal agonistic acts (i.e., fights) also differed according to the social status, the previous experience in the group and the observation day. Only fights initiated on days 0 and 2 were considered in the analyses since the incidence of fights on days 27 and 29 was minimal. Avoiders were not considered in those analyses since, by definition, they were not involved in fighting. First of all, the frequency of fights in which sows were involved strongly decreased from 0.88 [0.81;0.95] to 0.15 [0.13;0.18] fights / hour between day 0 and day 2 (*F*_1,939_ = 410.6, *P* < 0.0001). Dominant and Subdominant sows initiated more fights (0.26 [0.22;0.31], 0.25 [0.21;0.29] vs. 0.08 [0.06;0.10] fights / hour, *F*_2,939_ = 37.8, *P* < 0.0001) and received more fights (0.24 [0.20;0.28], 0.24 [0.20;0.28] vs. 0.10 [0.08;0.13] fights / hour, *F*_2,939_ = 23.4, *P* < 0.0001) than Loser sows between day 0 and day 2, independently of the observation day. In total, Dominants and Subdominants were involved in more fights per hour than Losers (**Fig 4a**, *F*_2,939_ = 62.0, *P* < 0.0001). The previous experience in the group also had an impact on fight frequency and New sows were more likely to be involved in fights than Resident sows (**Fig 4b**, *F*_1,939_ = 25.7, *P* < 0.0001).

**Fig 4.**
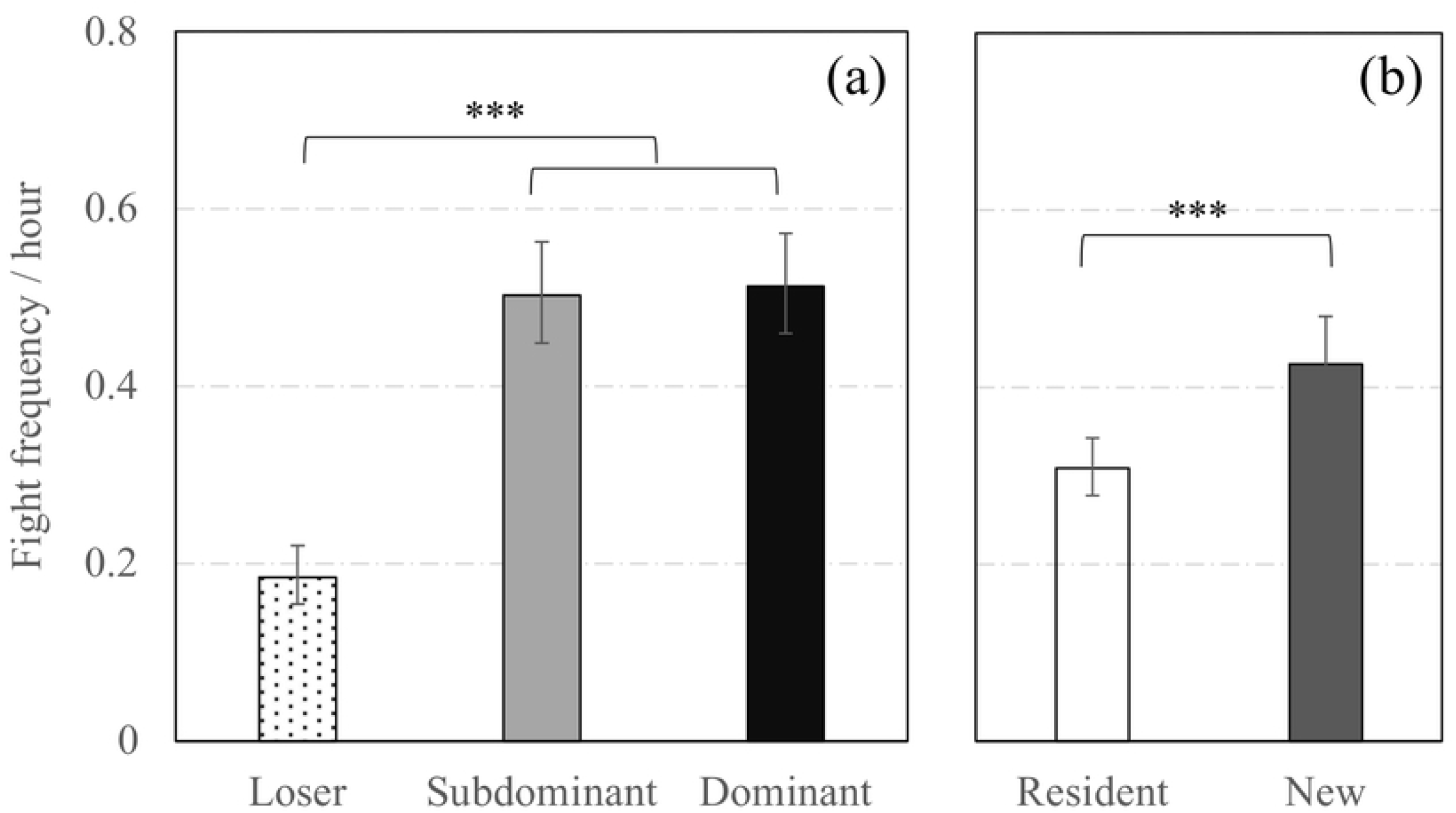
Fights around mixing according to social status and previous experience in the group. Back-transformed predicted mean [CI] number of fights per hour independently of the observation day (i.e., both day 0 (mixing day) and day 2) according to (a) the social status and (b) the previous experience in the group. ***; Means significantly differ (*P* < 0.001).

There was a triple interaction between the social status, previous experience in the group and observation day on the mean time spent fighting per hour (**Fig 5**, *F*_7,1119_ = 5.7, *P* < 0.0001). The longest time spent fighting concerned the Dominant sows newly introduced in groups (i.e., New Dominant) at mixing while both Resident and New Loser sows were the least involved in fighting. The **Fig 6** indicates that mean fight duration was longer for Dominants and Subdominants than for Loser sows at mixing but decreased and did not differ between social statuses on day 2 (*F*_2,532.4_ = 5.3, *P* = 0.005). Finally, fights lasted an average of 13.03 [11.30;15.03] sec in New sows while they lasted an average of 11.30 [10.10;12.65] sec in Resident sows (*F*_1,758.4_ = 3.1, *P* = 0.075).

**Fig 5.**
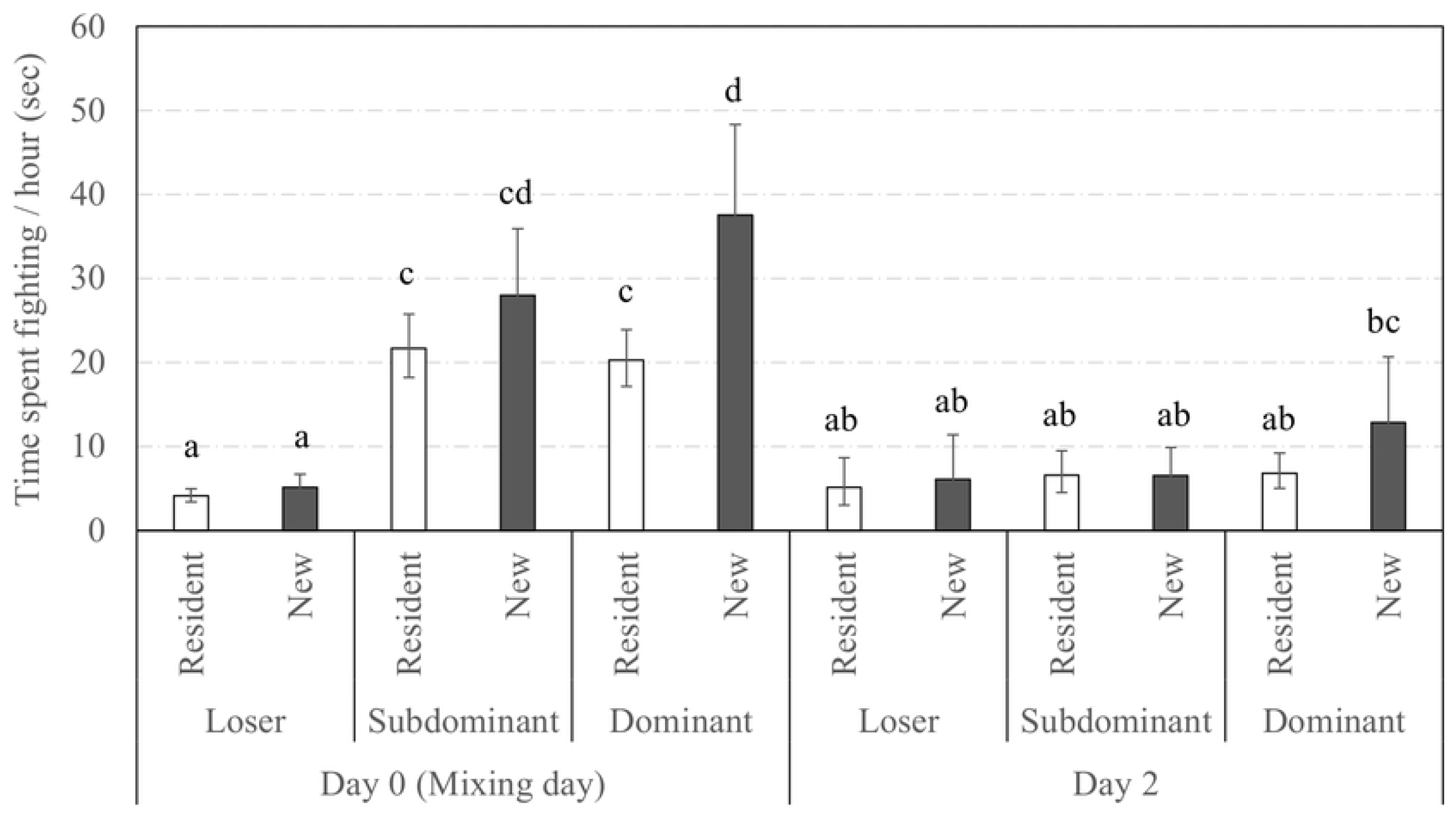
Time spent fighting across days according to social status and previous experience in the group. Back-transformed predicted mean [CI] time spent fighting (sec) per hour across observation days and according to the social status and the previous experience in the group. Different superscript letters indicate categories that significantly differ (*P* < 0.050).

**Fig 6.**
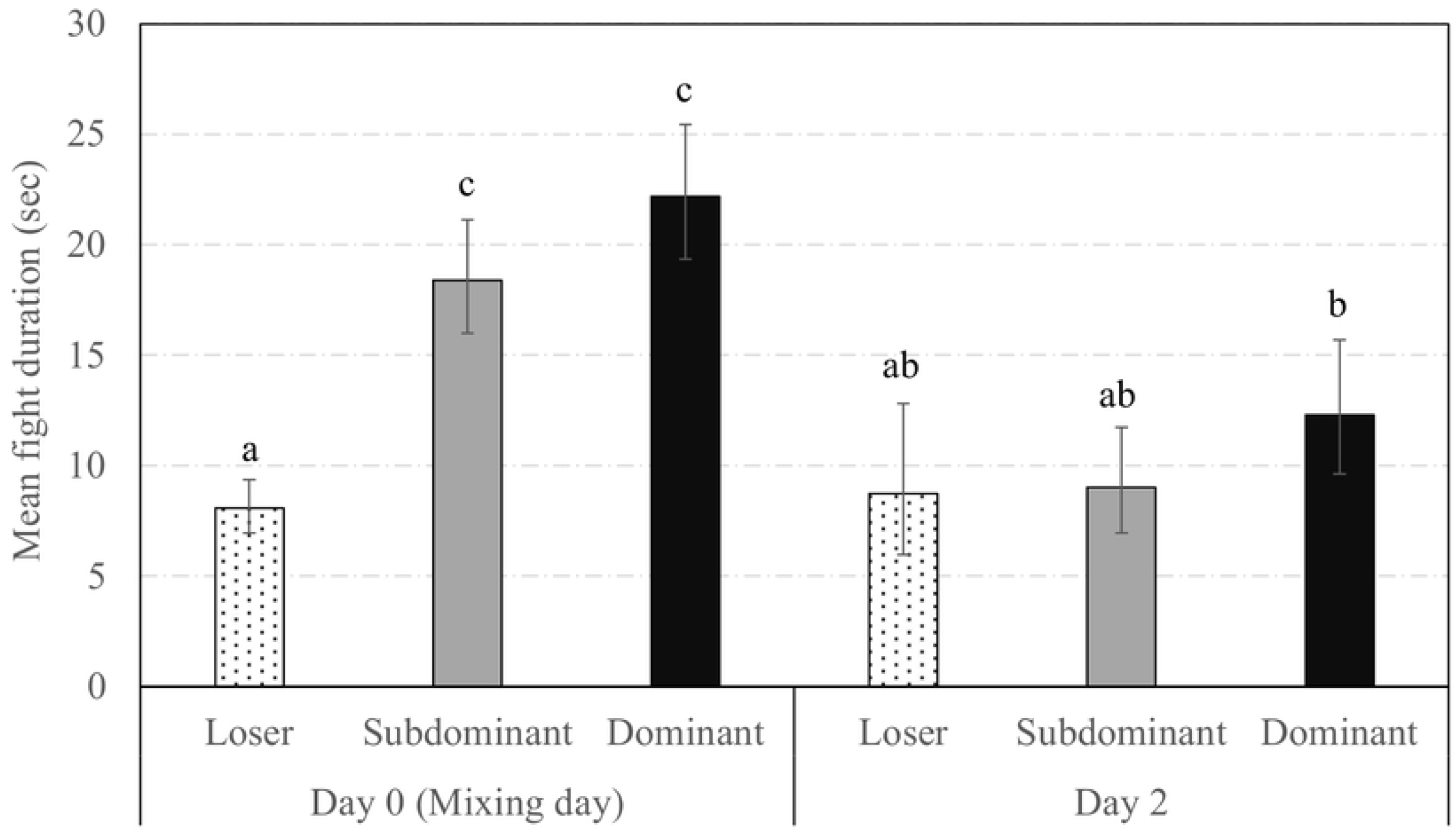
Fight duration across days according to social status. Back-transformed predicted mean [CI] fight duration (sec) across observation days and according to the social status. Different superscript letters indicate categories that significantly differ (*P* < 0.050).

### Body lesions

The social status had a significant impact on body lesion scores that differed according to the day post-mixing (**Table 2**). On the day after mixing, the highest total body lesion scores were seen in Subdominant sows and then in Dominant sows. Losers had significantly lower body lesion scores than high-ranking sows (Losers vs. Dominants: OR = 0.50, *t*_1,2076_ = −3.7, *P* = 0.014; Losers vs. Subdominants: OR = 0.27, *t*_1,2076_ = −6.6, *P* < 0.0001). However, this is Avoiders who had the lowest body lesion scores, even lower than Losers (OR = 0.33, *t*_1,2076_ = −5.9, *P* < 0.0001), confirming that they could successfully avoid aggression around mixing.

**Table 2.** Summary of the impact of social status on body lesion scores on each day post-mixing.

| Body region | Social Status | d1 |  | d26 |  | d84 |  |
| --- | --- | --- | --- | --- | --- | --- | --- |
|  |  | OR<sup>a</sup> | *P* | OR<sup>a</sup> | *P* | OR<sup>a</sup> | *P* |
| Front | Subdominant | 1.23 | ns | 1.59 | **0.025** | 1.08 | ns |
|  | Loser | 0.46 | **0.003** | 1.90 | **0.0004** | 1.80 | **0.009** |
|  | Avoider | 0.12 | **<0.0001** | 2.21 | **<0.0001** | 2.08 | **<0.0001** |
| Middle | Subdominant | 1.99 | **0.020** | 1.82 | **0.012** | 1.37 | ns |
|  | Loser | 0.74 | ns | 2.07 | **0.0004** | 1.99 | **0.011** |
|  | Avoider | 0.36 | **<0.0001** | 2.98 | **<0.0001** | 2.61 | **<0.0001** |
| Rear | Subdominant | 1.53 | ns | 1.28 | ns | 0.97 | ns |
|  | Loser | 0.66 | ns | 1.33 | ns | 1.10 | ns |
|  | Avoider | 0.39 | **0.0004** | 1.86 | **<0.0001** | 1.50 | ns |
| Total | Subdominant | 1.83 | **0.048** | 1.79 | **0.013** | 1.17 | ns |
|  | Loser | 0.50 | **0.014** | 2.31 | **<0.0001** | 1.76 | *0.070* |
|  | Avoider | 0.16 | **<0.0001** | 2.99 | **<0.0001** | 2.36 | **<0.0001** |

<sup>a</sup> The reference level is the Dominant status thus odd ratios higher than 1.00 indicate that lesion scores of sows from other social status (i.e., Subdominant, Loser or Avoider) are higher than lesion scores of Dominants.

Dominant sows had the lowest scores later in gestation, on d26 and d84 post-mixing while Avoider and Loser sows had the highest. At this time, lesion scores of Avoider and Loser sows did not differ significantly from one another (*P* > 0.10). Subdominant sows had an intermediate position. Indeed, Avoiders tended to have higher body lesion scores than Subdominants on d26 (OR = 1.67, *t*_1,2076_ = 3.2, *P* = 0.067) and they had significantly higher scores on d84 (OR = 2.03, *t*_1,2076_ = 3.9, *P* = 0.005). However, Subdominants did not significantly differ from Losers either on d26 or on d84 (*P* > 0. 10).

The previous experience in the group also had an impact on body lesions. Overall, New sows had more total body lesions than Resident sows, independently of the gestation phase (OR = 1.56, F_1,2076_ = 21.23, *P* < 0.0001).

### Feeding order (FO) at the ESF station

The feeding order (FO) between one-, two- and three-months post-mixing was highly consistent (0.79 < *r* < 0.86; *P* < 0.0001). Thus, the median FO between one- and three-months post-mixing was selected and used for subsequent analysis. Analyses indicated a strong association between the FO and social status (CMH: χ^2^ (6, N = 1121) = 74.22, *P* < 0.0001) with Dominant sows that had prior access to feed and Avoiders that were the last in the queue to feed. There was also a strong association between the FO and the previous experience in the group (CMH: χ^2^ (2, N = 1118) = 60.13, *P* < 0.0001) with more Resident sows that had prior access to feed over New sows.

### Individual and reproductive performance

Sow body weight before farrowing differed according to social status (*F*_3,179_ = 3.8; *P* = 0.011), with Dominant sows significantly heavier than Avoider sows (267.74 ± 1.95 kg vs. 259.31 ± 2.08 kg, respectively). Subdominant and Loser sows had an intermediate body weight and did not differ from both Dominant and Avoider sows (*P* > 0.10). There was no evidence of an impact of social status on any other performance measure (*P* > 0.10).

## Discussion

Animal welfare is about, above all, the extent to which an individual perceives and copes with its environment (15,16), rather than the population coping. Hence, the welfare level of individuals within a social group may differ from the welfare level of its other members, depending on several factors such as the social status (18,39). A previous study indicated that behaviour and welfare differed between groups according to the genetic line (32). In the present study, the individual social status of sows housed in large semi-static groups was determined based on the fight outcomes and its effect on the welfare of individuals within groups was tested. The originality of this study is that experiments were performed in a typical commercial farm setting with large groups up to 91 animals, compared to previous studies where the role of social status was generally explored in groups from several individuals to few dozen of individuals. Previous experience in the group (i.e., Resident or New) was also considered in the analyses which allowed to detect complementary findings.

### Social status classification and group composition

Dominance hierarchies yielded by contest outcomes structure pig populations, as observed in other gregarious species in the animal kingdom (40). The social status classification developed in this study was based on fight outcomes (i.e., proportion of fights won) around mixing (data from day 0 and day 2 post-mixing pooled) and was divided in three categories: the Dominants that won more fights than they lost, the Subdominants that lost more fights than the won, and the Subordinates that never won any fight. This latter category was then sub-divided in two sub-categories: the Losers that lost all the fights, and the Avoiders that were never involved in fights during the observation phases. Various social rank calculation methods have been used to categorise sows. While several authors attributed social ranks to sows based on displacement success at the feeding station (18,26) or fights success at mixing (20,27), other authors categorised animals according to their aggressiveness, either around mixing (23,41) or during a feed competition test (17,39). Although winning experience modifies individual’s own estimate of fighting ability and motivation to initiate a fight (42,43), in the present study, sows that were systematically aggressed or predominantly aggressors or with an intermediate behavioural pattern could be found in each category of Dominant, Subdominant as well as Loser, even if not in the same proportion. The percentage of agonistic acts won has already been positively correlated with displacement success at the feeding station (26) but it was decided to exclude displacement behaviours since those behaviours might be difficult to detect and may not be reliable indicators in large groups with high density of animals, especially around mixing when there is agitation in the pen. Aggression was also limited at feeding since sows were fed using non-competitive ESF systems (i.e., offering full protection to the sow at feeding, (35)). The authors are thus confident that fight outcomes around mixing was the best indicator of true dominance compared to other variables, including the propensity to initiate a fight that may also be influenced by the aggressive temperament or an internal motivation to dominate.

Sows in this study were observed around mixing (i.e., 4-hr at mixing and 2-hr at day 2 post-mixing), including just after mixing when contests are the most intense to establish hierarchy (44). At that time, subordinate sows (i.e., that never won any fight) represented almost half of the animals (i.e., 45.7 %). Among the subordinate sows, half of them were able to avoid fighting around mixing (i.e., the Avoiders). Also, agonistic interactions at the individual level seemed less frequent in this study compared to other studies with smaller group size. For instance, when focusing on the total frequency of agonistic acts (reciprocal and non-reciprocal) during the first two hours post-mixing, Dominant and Subdominant sows were engaged in approximately 4.9 to 6.8 agonistic acts per hour while Loser and Avoider sows were engaged in approximately 2.1 to 3.5 agonistic acts per hour. In groups of 6 or 26 multiparous sows, Li et al. (20) recorded rates of agonistic acts (i.e., defined as pushing and biting, but threats were excluded) almost twice as high during the first two hours post-mixing and those rates differed according to social status (i.e., based on winning success at mixing) and group size. Indeed, high-ranking sows were involved in approximately 9.5 (i.e., groups of 26 sows) or 11.5 (i.e., groups of 6 sows) agonistic acts per hour and low-ranking sows in approximately 4.5 agonistic acts per hour (i.e., both group sizes). Other factors than group size may explain differences between studies such as the individual floor space allowance (i.e., 2.21 m^2^/sow or more in this study vs. 1.50 m^2^/sow) or sows’ experience and age (i.e., mean parity of 3.2 in this study vs. 2.5). However, social strategies and organisation also differ according to group size and sows may adopt less aggressive social strategies in large groups, as previously theorized and demonstrated in weaned pigs by Andersen et al. (31). Indeed, the more the number of sows increases, the more the number of dominance relationships increases and the more a sow is likely to be severely injured. The adoption of a low aggressive strategy may reduce the risk and cost of being engaged in aggressive encounters. In addition, sows housed in large groups, and thus in larger pens, may have more ability to avoid an opponent. In the line with this hypothesis, Jensen (45) argued that aggression regulation within groups depends on the ability of subordinate sows to avoid the dominants rather than on the motivation of dominants to attack the subordinates. Social status is often divided in two (e.g., dominant and submissive pigs in Rhim et al. (46)) or three (e.g., low, middle or high social status in Zhao et al. (47)) categories and rare are the studies that divided animals in more than three categories (but see Greenwood et al. (36) and Turner et al. (48)). However, investigating dominance in a large group (up to 91 animals) resulted in the presence of a different category of sows (compared to previous studies with small groups), i.e., a quarter of the sows that did not participate in fighting at all during the observation phase (i.e., the Avoiders). In fact, the four social statuses highlighted in this study were balanced within groups since Dominants, Subdominants, Losers and Avoiders represented 29 %, 25 %, 23 % and 23 %, respectively.

Investigating social aspects on the sows’ welfare in a commercial setting of semi-static group management offered the opportunity to integrate the previous experience in the group factor since sows with gestation failures were re-inseminated and transferred to the next cohort. In total, groups were composed on the average of 69.58 ± 2.90 % Resident sows, defined as sows from the same group during the previous gestation, and an average of 30.42 ± 2.90 % New sows, defined as sows newly introduced into the group at mixing. Interestingly, preliminary analyses indicated that social status was not modulated by the previous experience in the group and both factors were thus considered in the analyses as complementary effects.

### Social status and welfare

Social status is an important factor for modulating health and welfare of the individuals, both at hierarchy formation and later in stable groups. Vying with an opponent to assert dominance often results in violent contests in pigs and is costly, but it arises because it offers stability and social cohesion to the group members and confers advantages to victorious animals on the long-term. However, this costly and risky behaviour may be avoided by weaker or previously defeated low-ranking animals that are less likely to be victorious. As expected, aggression behaviour was modulated by the social status, but this differed according to the observation day. This section is thus dedicated to the impact of social status on agonistic behaviours manifested across observation days. Triple interaction effects between social status, previous experience in the group and observation day were found when considering non-reciprocal agonistic acts delivered as well as total time spent fighting around mixing. Overall, the direction of social status x observation day interactions on behaviours was comparable between Resident and New sows, but behavioural differences between social statuses were numerically greater in New than in Resident sows. In addition, social status and previous experience in the group had dissociated effects on other behaviours (i.e., non-reciprocal agonistic acts received and fight duration) and body lesion scores. Hence, this section will mainly focus on the effects of social status. The effect of previous experience in the group x social status interaction will be addressed only when particularly relevant, but the impact of previous experience in the group on the welfare of sows will be addressed in the next section.

Avoider sows, that successfully avoided fights around mixing, were less likely to deliver non-reciprocal agonistic acts (i.e., bites, knocks, threats and bullying) and they were also bullied less than Losers and Subdominants. Their lesion scores recorded on the front, middle, rear and total body were the lowest on the day after mixing, confirming that they were successful at avoiding conflicts around mixing. This result contrasts with previous findings from Borberg and Hoy (21) who found a higher prevalence of skin injuries in subordinate sows after mixing in groups of 8 individuals. While subordinate sows may be the main targets of aggression from higher-ranking sows in small groups, they may be able to isolate themselves from opponents in larger groups where more space is available. However, although Avoiders’ welfare was favourable around mixing, they suffered from more aggression later during gestation. Indeed, it is interesting to note that Avoiders were eventually the main recipients of non-reciprocal agonistic acts one-month later, and they also had the highest body lesion scores one- and three-months post-mixing, as previously found in smaller group size (17). Feeding order (FO) was also associated with social status, as previously demonstrated (22,23) and Avoider sows were the last in the feeder queue while Subdominants and the Dominants had priority access to feed. Subordinate sows are more likely to be subjected to high levels of stress (49), but they may be able to cope by retreating to safe and less preferred areas (50,51) and feeding once Dominants are already fed to avoid conflicts. Avoider sows were characterized by a lower body weight before farrowing than Dominant sows and this body condition asymmetry may explain why those low-ranking sows avoided any fight around mixing. Alternately, Kranendonk et al. (26) claimed that high-ranking status sows may gain more body weight during gestation than low-ranking status sows due to their ability to displace low-ranking sows and steal little amounts of feed from the feeder. However, the ESF used in this study offered a full protection to the sows and prevented higher-ranking sows from displacing conspecifics while feeding.

Loser sows were not significantly lighter than Subdominant and Dominant sows but they still lost all the contests in which they were involved around mixing. They gave more non-reciprocal acts than Avoiders but less than higher-ranking sows at mixing and still had an intermediate position on day 2. Also, they were among the principal recipients of non-reciprocal agonistic acts on day 2. They had less total body lesions than higher-ranking sows around mixing, more likely because they were less involved in harmful reciprocal fights both in terms of number and total duration of fights (37). Regardless of their aggressive pattern (i.e., 52 % of them never initiate a fight, but 31 % were predominantly aggressors), Losers may rapidly evaluate their weakness compared to their opponents since the mean fight duration was lower at mixing compared to their higher-ranking counterparts. At day 29, they still gave numerically less non-reciprocal agonistic acts and received numerically more non-reciprocal agonistic acts than higher-ranking sows. In terms of body lesions, they had more total body lesions one-month post-mixing and they tended to have more three-month post-mixing than Dominant sows, but they did not differ from Subdominants and Avoiders both one- and three-months post-mixing.

Subdominant sows were the ones that had the most affected welfare state, in terms of stress, aggression and body lesions, during hierarchy formation. The establishment of dominance hierarchy is usually particularly detrimental for Subdominant sows that are highly involved in fighting to assert their dominance but lose a high number of fights against their Dominant opponents. Like Dominant sows, Subdominants were characterized by high levels of aggressiveness since they gave a high proportion of non-reciprocal agonistic acts and fights around mixing. They were also among the main recipients of non-reciprocal and reciprocal aggression around mixing. Overall, Subdominants had significantly more total body lesions after mixing than sows from higher or lower social rank. There is evidence that pigs highly involved in reciprocal fights have more skin lesions immediately after mixing, but likewise the strong propensity to receive non-reciprocal aggression is susceptible to increase body lesion severity (25). Once established, stable dominance hierarchy among group members can suppress unnecessary fights (52), which are among the most damaging social interactions, and future agonistic interactions are usually limited to single bites, knocks and threats due to the competition for resource access (e.g., feeding, resting area), hunger, discomfort, irritability or frustration (8,53,54). On days 27 and 29, Resident Subdominants gave numerically more non-reciprocal agonistic acts than their Resident counterparts. Because sows were not frequently involved in fighting anymore, Subdominant sows, but also Dominants, were simply rarely recipients of agonistic interactions and this was confirmed by body lesion scores that were significantly lower than those of Avoiders, one- and three-months post-mixing.

Asserting dominance is a costly behaviour which exists only because it confers advantages to dominant animals, such as prior access to resources. For example, dominant sows were observed more frequently in prime lying areas than subordinate sows in the study of O’Connell et al. (23). Dominants, which were characterized by a higher body weight before farrowing than Avoider sows, indeed had priority access to the feeder. Although winning experience may modify an individual’s estimate of fighting ability and motivate to initiate a fight, Dominant individuals are not necessarily always the most aggressive. They did not differ from Subdominants in their propensity to initiate or to receive a fight, or in fight duration. However, they gave significantly more non-reciprocal agonistic acts while they received less around mixing. As a result, they had intermediate body lesion scores at mixing since their skin was more injured than that of Losers and Avoiders, but less than that of Subdominants. One-month later, on day 29, Resident Dominant sows were not the most aggressive among the Resident sows while the New Dominant sows were numerically the most aggressive among the New sows. However, Dominant sows received significantly fewer bites and knocks, threats and bullying than their lower-ranking counterparts, independently of the previous experience in the group. Overall, Dominants had the lowest body lesion scores one- and three-months post-mixing and this corroborates results from Tönepöhl et al. (25) that indicated the involvement in reciprocal fights around mixing was accompanied by lower skin lesions later during gestation.

Verdon et al. (55) investigated whether forming groups of aggressive sows would predict the subsequent aggression at mixing and they found no effect since groups formed with selected aggressive sows were not more aggressive at mixing than control groups. They argued that individuals may show behavioural flexibility and their aggressive predisposition may be dependent on the pen mates’ characteristics. In this study, the sows’ social status was fairly consistent across the two first replicates in which individuals were involved, suggesting that group characteristics may be comparable across replicates since 70% of the group members were the same across replicates (i.e., the Resident sows). In the commercial context in which this study was performed, New pigs are mixed with Resident pigs and a certain level of aggression around mixing is necessary. Dominant individuals, comparatively to the other pen mates, may be determinant for establishing dominance relationships and ensuring future social cohesion (56). Although Dominant and Subdominant animals may suffer from overt aggression around mixing, they may benefit from better welfare later in gestation compared to Subordinate animals (i.e., Losers and Avoiders in this study) that may pay the cost of being lower in hierarchy. Aggression in large groups appears to be less severe around mixing since sows may adopt less aggressive social strategies and some of them (i.e., the Avoiders) may be able to isolate themselves from aggressive opponents and avoid conflicts. However, low levels of aggression during the stressful event of mixing may solely delay the conflicting period (6), and social cohesion might be looser in large groups.

### Previous experience in the group and welfare

Resident sows benefited from their position in the group since they had priority access to the ESF compared to New sows once the hierarchy was formed. Agonistic behaviours and skin injuries were also determined by the previous experience in the group. Aggression has been shown to be exacerbated among sows that are unfamiliar with one another (57) and in this study, New sows were the main perpetrators of fights. Also, mean fight duration of New sows tended to be longer than that of Resident sows, indicating that contest resolution might be harder when sows are newly introduced in groups than when they were already resident in the group during the previous gestation. Indeed, New sows would have been much less likely to encounter sows they had previously fought, compared to Resident sows who were already part of the same group and represented the highest proportion of the group members (i.e., 70%). Regardless of the observation day, New sows had the highest body lesion scores highlighting the damaging effect of introducing new sows into groups to sows’ welfare. Hence, mixing animals that have already become known to each other in the previous gestation can shorten the time it takes to reorganise the social structure and preserve welfare. New sows, especially with a low social status, were more vulnerable throughout gestation and could serve as indicators of non-optimal conditions. Indeed, farm personnel could detect vulnerable sows using the ESF monitoring software (i.e., vulnerable sows are the last to feed) and then pay more attention to their welfare state during health routine visits, by supposing that vulnerable sows with good welfare status may indicate that group welfare is acceptable.

The fact that Resident sows fought less around mixing (i.e., in terms of fight frequency and mean fight duration) indicates that pigs may have good social recognition skills. Even juvenile pigs can discriminate equally familiar conspecifics (58) and they may use their knowledge to adjust their foraging tactics to whom they are with (59). However, the extent to which individual recognition of all group members is plausible in groups of 50 or more individuals, as is the case in sheep (60), is unknown. Fraser and Broom (61) estimated that pigs may recognise between 20 and 30 counterparts and Verdon and Rault (62) suggested that individual recognition might be less likely in unnaturally large group sizes. Although it is not possible to know whether Resident sows recognised (*“Do I know you?”*) all other Resident group members and had a mental representation of individuals’ identity using this experimental design, this study presupposes that they are, at least, able to discriminate (*“Are you different?”*) between resident and new individuals, as already demonstrated in growing pigs by Turner et al. (30).

Interestingly, New Dominant sows were highly involved in fights and they spent significantly more time fighting per hour than Resident Dominant sows, while this effect was not observed between lower-ranking sows. This finding suggests that New Dominant sows may have to get involved in a higher number of fights *to get known* by opponents and assert their dominance in their new social group whereas Resident Dominant sows may not have to rebuild their social network, and simple *reminders* to previous opponents or fights with new group members only might be sufficient to reassert their dominance. Compared to Dominant sows, there was no difference between Resident and New sows from Loser and Subdominant status. Rather than a lack of social memory, those sows may have to get involved in fights to improve or assert their social status, independently of their familiarity level (i.e., Resident vs. New).

## Conclusion

The welfare of sows differed between individuals within the social groups according to their social status, based on the fight winning, and their previous experience in the group (i.e., whether they were already Resident in the group in the previous gestation or New). This study draws attention to the social strategies adopted in large groups (up to 91 animals), as sometimes encountered in pig industry, that differ from the ones used in small groups. Here, a quarter of sows were able to avoid fighting around mixing (i.e., the Avoiders), suggesting that subordinate sows are more likely to avoid aggressive individuals when mixed in large pens with higher group size. As a result, those subordinate sows had the lowest lesion scores around mixing. However, subordinate sows (i.e., the Avoiders and Losers) suffered from more body lesions than higher-ranking sows later during gestation. In addition, this study showed that sows newly introduced into the groups paid the cost of being new since they were more involved in agonistic interactions and maintained higher body lesions scores, not only at mixing, but also later during gestation. Many questions remain open and future studies may thus investigate more deeply the social relationships of sows housed in large groups, including the extent to which pigs can develop preferred affiliative relationships with specific partners, as this is the case in growing pigs housed in small groups (63), and become part of a cohesive social group. The popularisation of the automatic detection of behaviours (64–68) will help to gather large amounts of data to study social relationships within large groups of sows.

## Acknowledgements

The authors wish to thank the piggery staff and Agri-Marché team for their valuable support during the experiments and the animal care. The authors are grateful to Dr. Sabine Conte for helping with the protocol development as well as Jean-Michel Beaudoin, Lucie Galiot and Béatrice Hamel for their participation in the body lesion scoring.

